# Spatial and semantic memory reorganize a hippocampal long-axis gradient

**DOI:** 10.1101/2025.10.15.682651

**Authors:** Anikka G. Jordan, Joel L. Voss, James E. Kragel

**Affiliations:** Department of Neurology, University of Chicago, Chicago IL 60637, USA

## Abstract

The hippocampus supports episodic memory by binding spatial and semantic information, yet how this information is simultaneously organized along its long axis remains debated. Gradient accounts propose a continuous shift in representational scale, from coarse coding in anterior to fine coding in posterior regions, whereas modular accounts posit discrete subregions specialized for distinct functions. Using high-resolution fMRI together with eye tracking as a readout of spatial and semantic memory during sequence learning, we directly tested these competing models. During predictable sequences, hippocampal activity continuously varied along the long axis. In contrast, modular organization emerged when sequences mismatched memory. Subregions in the anterior and posterior hippocampus were sensitive to semantic and spatial mismatches, respectively. Notably, the intermediate hippocampus was specifically sensitive to concurrent mismatches in both dimensions, but not to mismatches in either dimension alone. These content-sensitive subregions were embedded within distinct cortical networks that reorganized according to memory demands. Together, our findings show the hippocampus flexibly combines gradient and modular dynamics to simultaneously represent the spatial and semantic content that defines episodic memory.

## Introduction

The hippocampus supports episodic memory [1–3] with neuronal activity containing spatial [4, 5] and semantic [6, 7] information. While the hippocampus is often viewed as a single homogeneous structure, research has revealed differences in its functionality [8], structure [9, 10], and genomic characteristics [11] along its longitudinal axis. There is strong evidence for a gradient of spatial coding in rodents, as the receptive fields of place cells that code for the location of a rodent in the environment shrink in the ventral to dorsal direction [12]. Based on these findings, models of hippocampal organization have been proposed wherein there are gradients of function along the long axis [13]. Although there is evidence for differences in structure [14] and function [15] along the hippocampal long axis in humans, how this organization relates to the spatial and semantic content of memories remains to be determined.

In humans, the posterior hippocampus is theorized to process detailed, visuospatial information, whereas the anterior hippocampus is thought to support more abstract, meaning-based (i.e., semantic) representations [13, 16, 17]. Evidence for functional gradients is limited to a handful of studies that have examined visual [18], spatial [19], and abstract [20] representations in isolation. It therefore remains unclear how the hippocampus supports simultaneous processing of the spatial and semantic representations that define episodic memories. Gradient accounts propose that representations shift smoothly along the long axis, with the intermediate hippocampus reflecting a midpoint in representational scale [8]. Modular accounts suggest that anterior, intermediate, and posterior regions act as distinct functional modules [21], with the intermediate hippocampus potentially integrating activity at either end of the long-axis. Distinguishing between these gradient and modular accounts will therefore reveal how the hippocampus transforms spatial and semantic inputs into unified memory representations.

Differences in connectivity along the hippocampal long axis may explain how the anterior and posterior hippocampus support distinct functions. The posterior hippocampus is known to be part of the parietal memory network [22, 23]. The hippocampal tail is also associated with the salience network, although the precise contributions of this network to cognition have not yet been clearly defined [24, 25]. Recent fMRI studies have used connectopic mapping (also referred to as gradient mapping, 26) to characterize hippocampal organization based on changes in connectivity along its long axis [27]. The posterior hippocampus exhibits greater connectivity to early visual and dorsal occipito-parietal regions [28]. Meanwhile, the anterior hippocampus exhibits greater connectivity to regions in the default network, including the angular gyrus, anterior temporal lobe, and posteromedial cortex, which support self-oriented processing like autobiographical memory [28, 29]. These differences in connectivity suggest an alternative to gradient-based organization [19, 20], namely that distinct cortical networks anchor anterior and posterior hippocampal functions. A central theoretical question is therefore whether hippocampal long-axis organization reflects a continuous gradient of representational scale or modular specialization driven by network-level affiliations. Clarifying this distinction is essential for linking cellular models of hippocampal computation to systems-level accounts of human memory.

One major function of the hippocampus is to detect mismatches [30–32], which are prediction errors that occur when incoming sensory inputs conflict with stored relational memory structures. Hippocampal activity scales with the degree of such mismatches across a range of content types, including unexpected or out-of-sequence objects [33, 34], and violations of expected visual categories [35]. These findings suggest a general mechanism for detecting relational change [34]. Yet mismatches can arise from different types of representations, including spatial and semantic, and how these distinct representational demands shape functional organization along the hippocampal long axis remains unclear. This region thus offers a strong test of competing models. If gradient-based organization dominates, mismatch effects should scale smoothly along the long axis regardless of content. If modular accounts hold, the intermediate hippocampus may emerge as a critical locus where spatial and semantic prediction errors converge.

We therefore designed an fMRI experiment to characterize the organization of spatial and semantic information along the hippocampal long axis during a sequence memory task. By using eye tracking to quantify spatial and semantic memory behavior on the task with high temporal precision [36], we hoped to identify hippocampal mismatch signals reflected in memory-guided behaviors [37]. By including both spatial and semantic mismatches, we were able to test whether similar organizational principles hold for these distinct types of information. In addition, we tested the degree to which hippocampal long-axis organization could be explained by the connectivity of distinct cortical systems.

## Results

### Semantic and spatial mismatch strength enhances sequence discrimination

We developed an fMRI sequence-memory paradigm (Fig. 1A) to investigate how hippocampal mismatch signals for semantic and spatial information vary along its long axis (see Movies S1-S4 for example trials and gaze behavior). During the task, participants learned sequences of five objects presented in different locations on a circular array. Semantic and spatial transitions between objects in each sequence followed a predictable order. Semantic transitions were either near or far based on a multidimensional space of object features (38, see Methods). Spatial transitions were either 45 degrees (Near) or 180 *±* 45 degrees apart. These sequences allowed us to vary the magnitude of mismatch signals based on the similarity of spatial or semantic content at test, while maintaining familiarity for objects and locations within a given sequence.

**Figure 1.**
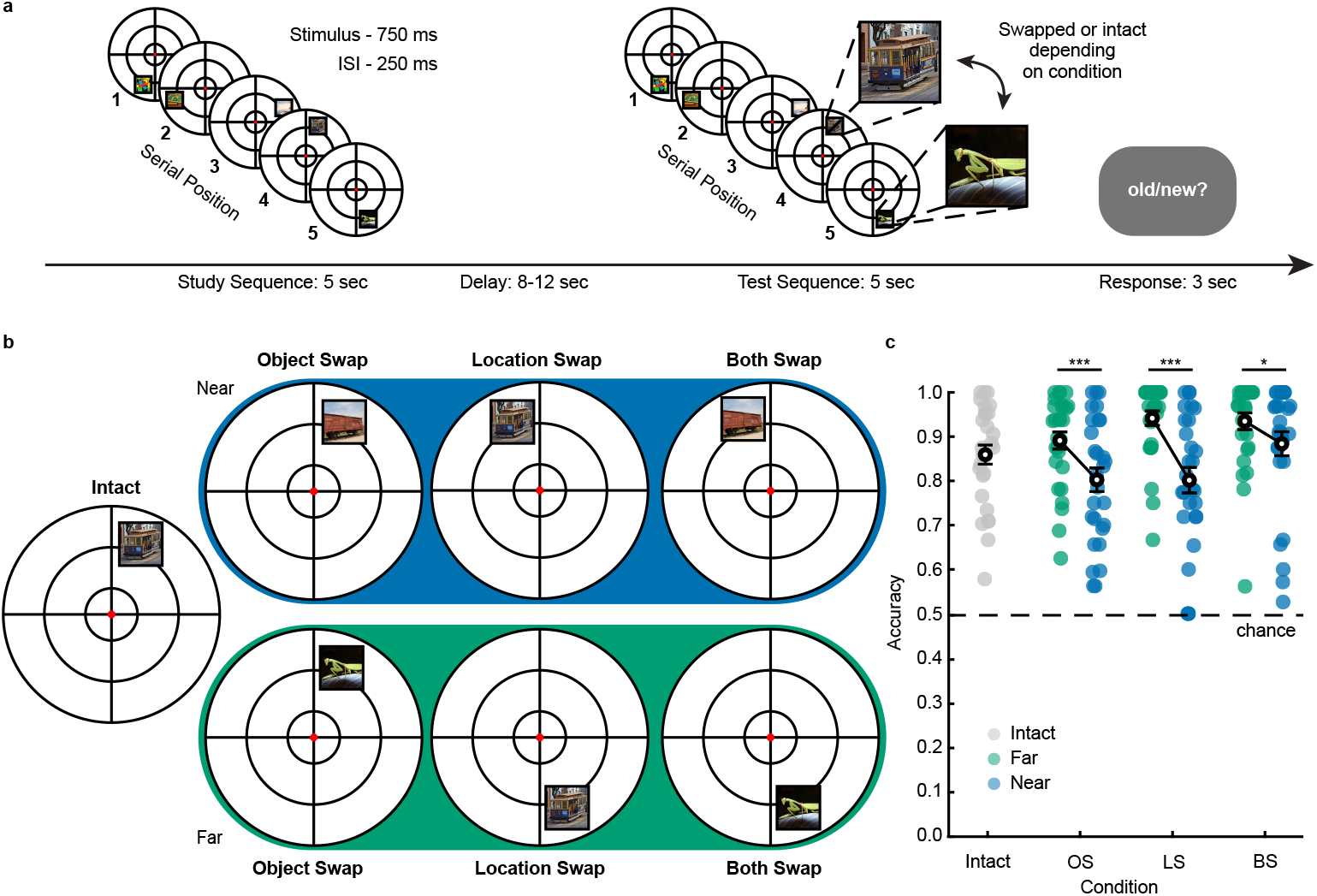
Spatial and semantic mismatches enhance sequence discrimination. (**a**) Example trial sequence. In each 5 s sequence, participants viewed five objects presented at eight possible locations on a circular grid. Each object was presented for 750 ms with a 250 ms inter-stimulus interval (ISI). Following a delay, participants viewed a test sequence and indicated whether the sequence was old or new. (**b**) Schematic showing possible test conditions for the critical fourth stimulus in the sequence in **a**. Participants determined novelty on intact (I), object swap (OS), location swap (LS), or both swap (BS) sequence when stimulus attributes were swapped between the last two serial positions in the sequence. Swap distances were either near or far based on the semantic (top, see Ref. 38) or spatial features of the stimulus. (**c**) Accuracy for each test condition. Participants (*N* = 28 participants, depicted by circles) performed significantly above chance (indicated by the dashed line) and were better at detecting far compared to near mismatches. Error bars denote SEM. ^*\*\*\**^, *p <* 0.001.

Following a brief delay, participants judged whether a test sequence matched the initial sequence. One fourth of the test sequences were intact. We manipulated the remaining sequences by swapping either the semantic, spatial, or both attributes of the final two stimuli in the sequence. Importantly, half of the swaps occurred on near and half on far transitions, varying the mismatch between the study and test sequences (Fig. 1B). Following the test probe, participants indicated whether each test sequence was old or new during a 3-second response period. This mismatch detection task allowed us to map both spatial and semantic hippocampal representations simultaneously.

Participants performed well on the task, with an overall accuracy of 87.1 *±* 1.7%. Pairwise comparisons of mismatch distance (near vs. far) revealed lower accuracy for near (82.9 *±* 2.2%) compared to far (92.3 *±* 1.5%) sequence manipulations (*F*_1,29.9_ = 31.3, *p <* 0.001; Fig. 1C). This indicates that the magnitude of the semantic and spatial mismatch influences task performance, with larger mismatches being easier to detect.

Examining each condition individually, participants were less accurate on near trials for object swap (*t*_27_ = 4.07, *p <* 0.001, *g* = 0.71 [0.33, 1.11]), location swap (*t*_27_ = 5.31, *p <* 0.001, *g* = 1.11 [0.64, 1.63]) and both swap (*t*_27_ = 2.52, *p* = 0.018, *g* = 0.40 [0.07, 0.75]) manipulations. These results further confirm that task performance scales with the magnitude of the mismatch for each type of sequence manipulation.

If different types of information integrate during mismatch computation, the largest mismatch should occur when both the object and location are swapped. Indeed, both swap trials had the highest overall accuracy (90.9 *±* 2.1%), which differed significantly from object swap trials (84.7 *±* 2.0%; *t*_27_ = 3.81, *p* = 0.001, *g* = 0.56 [0.24, 0.89]) but not location swap trials (87.2 *±* 1.9%; *t*_27_ = 1.87, *p* = 0.073, *g* = 0.35 [*−*0.03, 0.74]). Moreover, differences between both swap and location swap trials varied with the magnitude of the mismatch (*t*_27_ = 2.79, *p* = 0.010, *g* = 0.71 [0.19, 1.25]). Participants were more accurate on both swap trials than location swap trials for near (*t*_27_ = 2.59, *p* = 0.015, *g* = 0.55 [0.11, 1.00]) but not far (*t*_27_ = *−* 0.39, *p* = 0.702, *g* = *−* 0.07 [*−* 0.46, 0.31]) mismatches. These findings indicate that the paradigm is sensitive to the magnitude of semantic and spatial mismatches and their combination, supporting the idea that both types of information contribute to mismatch computation.

### Eye movement behavior predicts semantic and spatial mismatches

Previous research has demonstrated eye tracking can be used to look at memory-guided (predictive) eye movements [37, 39, 40]. If participants successfully learned the spatial sequence during study (Fig. 2A), they should be able to focus their attention and saccade to upcoming object locations prior to their appearance at test. We asked if spatial memory was reflected in eye movements by comparing the number of fixations to regions of interest on the visual array (Fig. 2B). Specifically, we asked whether participants were more likely to fixate on the original location from the study period as compared to other possible object locations in the array. We also measured spatial mismatches by quantifying the number of fixations to upcoming swapped locations.

**Figure 2.**
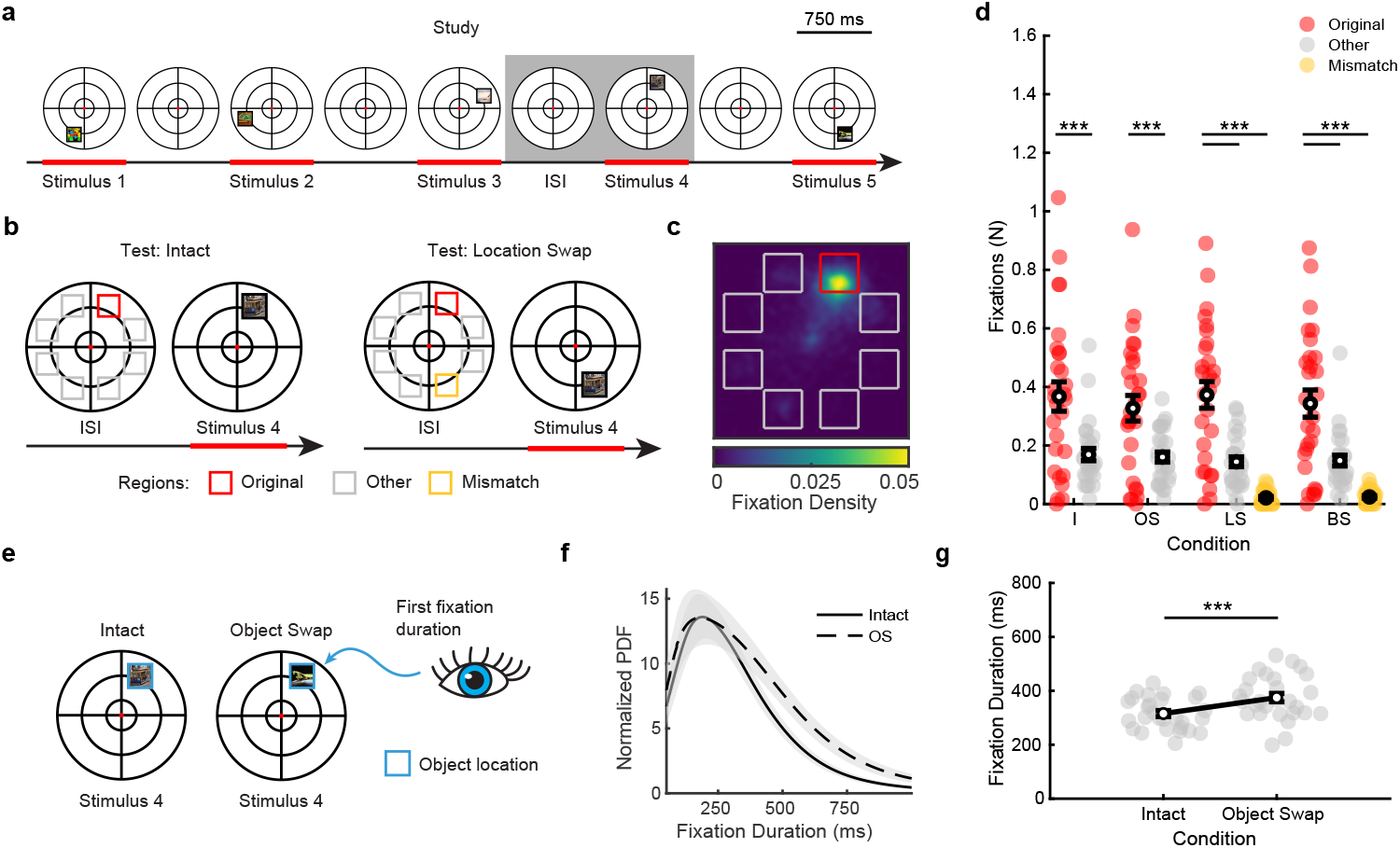
Eye movements reflect spatial and semantic memory. (**a**) Example study sequence with inter-stimulus intervals (ISIs). The grey box highlights the stimulus at serial position four and the preceding ISI, which are critical timepoints at test. (**b**) Schematic showing regions of interest to measure spatial memory. We measured fixation counts to the original location of the upcoming object, denoted by a red square. On location swap (LS) and both swap (BS) trials, we also measured fixations to the mismatched location (denoted by a yellow square). Other potential object locations (grey squares) served as controls. (**c**) Spatial heat map of all fixations during the ISI preceding stimulus four at test. Fixation maps were rotated to align the original object locations. Regions are colored as in **b**. (**d**) Effect of spatial memory on viewing behavior. Participants made significantly more fixations to the original than other object locations, reflecting spatial memory. On LS and BS trials, participants made more fixations to the original compared to mismatched object locations (*N* = 28 participants, depicted by circles; paired-samples t-test). (**e**) Schematic showing regions of interest to measure semantic memory. We measured the duration of the first fixation to the object in stimulus position four at test (blue square). (**f**) Probability density function of first fixation durations for intact (I) vs. object swap (OS) trials showing their different distributions. Error bars denote SEM. (**g**) First fixation durations increased for OS relative to I trials (paired-samples t-test). Error bars denote SEM. ^*\*\*\**^, *p <* 0.001.

As shown in Figure 2C, participants primarily fixated on the original study location in the moments prior to its presentation at test. Indeed, participants made more predictive than other fixations on intact (*t*_27_ = 5.42, *p <* 0.0001, *g* = 1.46 [0.84, 2.14]), object swap (*t*_27_ = 4.48, *p* = 0.0001, *g* = 1.34 [0.68, 2.06]), LS (*t*_27_ = 5.52, *p <* 0.0001, *g* = 1.60 [0.91, 2.35]), and both swap (*t*_27_ = 4.62, *p* = 0.0001, *g* = 1.29 [0.66, 1.98]) trials (Fig. 2D). We also confirmed this behavior produced spatial mismatches on both location swap (*t*_27_ = 7.47, *p <* 0.0001, *g* = 1.97 [1.26, 2.76]) and both swap (*t*_27_ = 5.66, *p <* 0.0001, *g* = 1.55 [0.89, 2.28]) trials when the location in memory mismatched the visual percept. Predictive fixations thus provide an online measure for spatial memory and indicate mismatch processing above and beyond recognition accuracy, which may reflect the outcome of deliberative processing of the entire test sequence.

Eye movement behavior also provides a sensitive measure of object memory, including objects within a spatiotemporal context [41–43]. Humans initially fixate for longer on unexpected compared to repeated [44, 45] or semantically typical [46] objects. Thus, we used first fixation durations to objects in the fourth sequence position (either mismatched or repeated) as a measure of object memory (Fig. 2E). At the critical swap period during test probes, participants initially fixated longer on unexpected relative to repeated objects (*t*_27_ = 5.17, *p <* 0.0001, *g* = 0.81 [0.46, 1.20]), reflecting their knowledge of the upcoming object (Fig. 2G). Increased viewing of unexpected objects may reflect memory for objects in the studied sequence, showing our task is sensitive to semantic memory.

### Semantic and spatial mismatches reorganize a common hippocampal gradient

To examine how neural representations are organized along the hippocampal long axis, we applied representational similarity analysis (RSA; 47) to multivoxel patterns within searchlights spanning the hippocampus (Fig. 3A). RSA quantifies the similarity of voxel patterns across trials and provides a proxy for how consistently local neuronal populations respond to different types of information. For example, if a region contains neurons tuned to spatial features, trials with similar spatial inputs should evoke similar patterns of activity [12, 18, 19]. In regions with narrower tuning, small differences in spatial input would lead to larger changes in neural similarity. To capture how these representational properties vary along the hippocampal long axis, we used a hippocampal unfolding algorithm that accounts for individual variability in morphology and enables alignment across participants [48–50].

**Figure 3.**
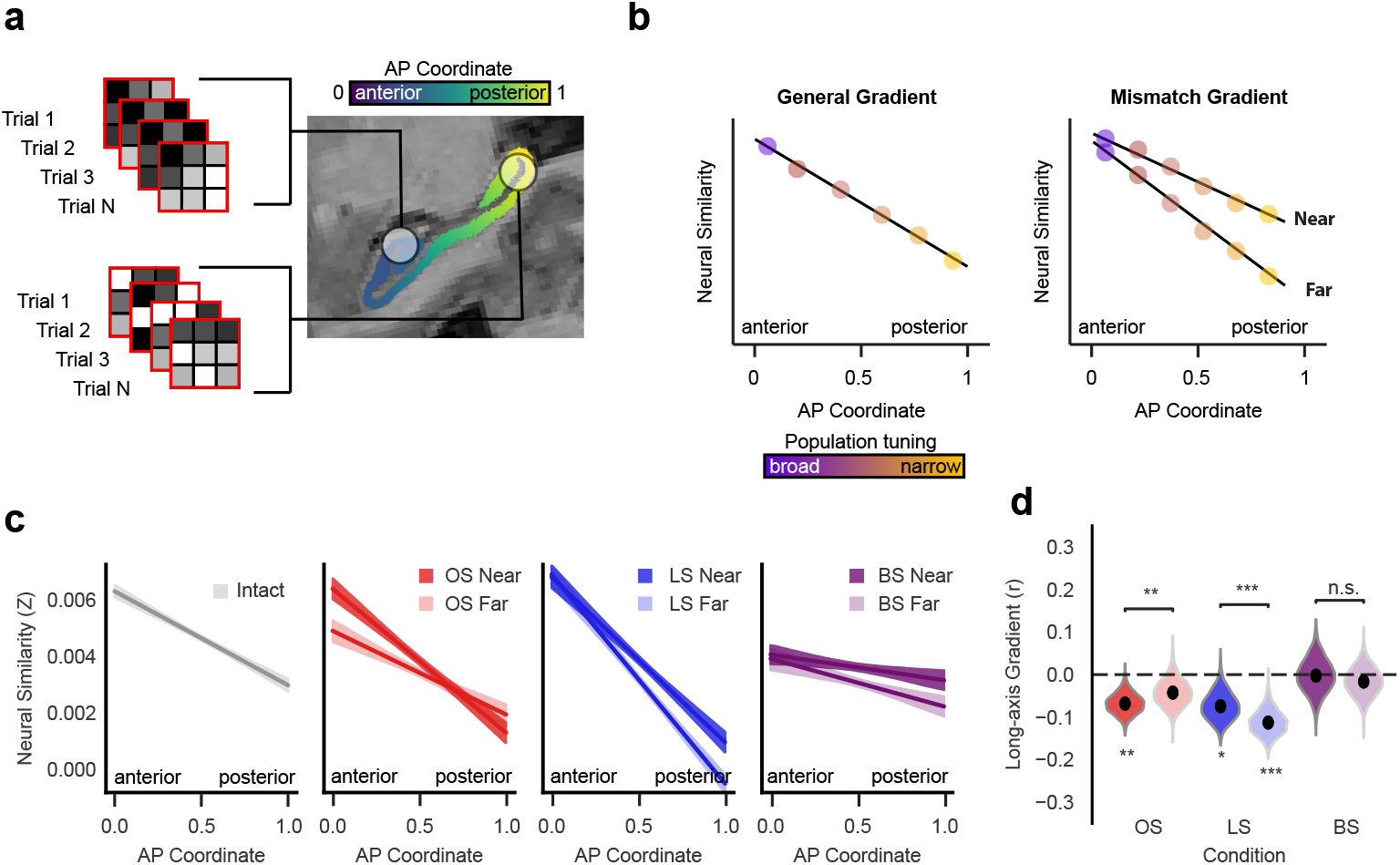
Mismatch processing disrupts a common hippocampal gradient. (**a**) Analysis schematic. Multivoxel patterns were extracted from hippocampal searchlights, and anterior–posterior (AP) gradients were tested. (**b**) Hypothesized neural similarity profiles. A general gradient would be unaffected by memory demands, whereas a mismatch gradient would scale with mismatch strength. (**c**) Group-level decreases in neural similarity along the AP axis (*N* = 28), with shaded 95% CIs. (**d**) Mismatch strength alters gradient slopes. Far object swaps reduce the AP gradient; far location swaps enhance it. Violin plots show bootstrap distributions of repeated-measures correlation coefficients; dots are subject means. ***, *p <* 0.001; **, *p <* 0.01; *, *p <* 0.05.

We then tested how representational organization differs as a function of content type (semantic versus spatial) and memory demands (degree of mismatch; Fig. 3B), considering two possible gradient-based organizational schemes. The first is a general gradient, in which neural similarity decreases from anterior to posterior hippocampus regardless of task condition, reflecting shifts in representational scale or tuning. The second is a mismatch-sensitive gradient, in which the strength of this anterior–posterior slope varies with mismatch magnitude, consistent with neural populations that encode the degree of change across events.

Under conditions where sensory input aligned with memory-based predictions, hippocampal activity exhibited a smooth anterior–posterior gradient, with maximal neural similarity in anterior hippocampus (aHPC; Fig. 3C). This gradient was most prominent during intact trials (*r* = *−*0.07, 95% *CI* [*−* 0.12 *−* 0.02], *p* = 0.006, permutation test) and when test sequences closely matched memory, including near object swaps (*r* = *−* 0.07, 95% *CI* [*−* 0.11 *−* 0.02], *p* = 0.007, permutation test) and near location swaps (*r* = −0.07, 95% *CI* [*−* 0.13 *−* 0.004], *p* = 0.02, permutation test), which did not significantly differ from repeats (OS: Δ*r* = *−* 0.003, 95% *CI* = [*−*0.015 0.015], *p* = 0.68, LS: Δ*r* = *−* 0.005, 95% *CI* = [*−* 0.016 0.016], *p* = 0.55, permutation test). Thus, when sensory inputs align with memory-based predictions, the hippocampus expresses a common long-axis gradient, regardless of whether the relevant content is spatial or semantic.

In contrast, large mismatches disrupted this organization. Far object swaps eliminated the anterior-posterior (AP) gradient (*r* = *−* 0.04, 95% *CI* = [*−* 0.10 0.02], *p* = 0.09, permutation test), producing significantly shallower slopes than near object swaps (Δ*r* = 0.02, *p* = 0.003) and intact sequences (Δ*r* = 0.02, *p* = 0.002). Conversely, far location swaps amplified the gradient, with steeper slopes than near location swaps (Δ*r* = *−* 0.04, *p <* 0.001) and intact sequences (Δ*r* = *−* 0.04, *p <* 0.001; Fig. 3C). When both semantic and spatial content mismatched memory, the AP gradient was absent for both near (*r* = *−* 0.004, 95% *CI* = [*−*0.08 0.07], *p* = 0.46, permutation test) and far (*r* = *−* 0.02, 95% *CI* = [*−* 0.08 0.04], *p* = 0.30, permutation test) swaps. These findings suggest that long-axis hippocampal organization dynamically reorganizes based on the strength and type of mismatch.

While these regression-based findings support a gradient-like functional architecture, they do not preclude the existence of discrete subregions atop this gradient. To identify such subregions, we compared neural similarity (at the searchlight center voxel) between near and far mismatches (Fig. 4 and Table S1). We identified four clusters with sensitivity to mismatch strength. Object swaps modulated neural similarity in left aHPC (*Z* = 2.86, *p <* 0.005); location swaps affected right posterior HPC (pHPC; *Z* = 3.05, *p <* 0.001); both swap types influenced similarity in left intermediate HPC (iHPC; *Z* = 3.11, *p <* 0.001) and right pHPC (*Z* = 3.04, *p <* 0.001). This spatially specific pattern mirrors the gradient findings, showing that semantic and spatial mismatches differentially engage regions along the long axis.

**Figure 4.**
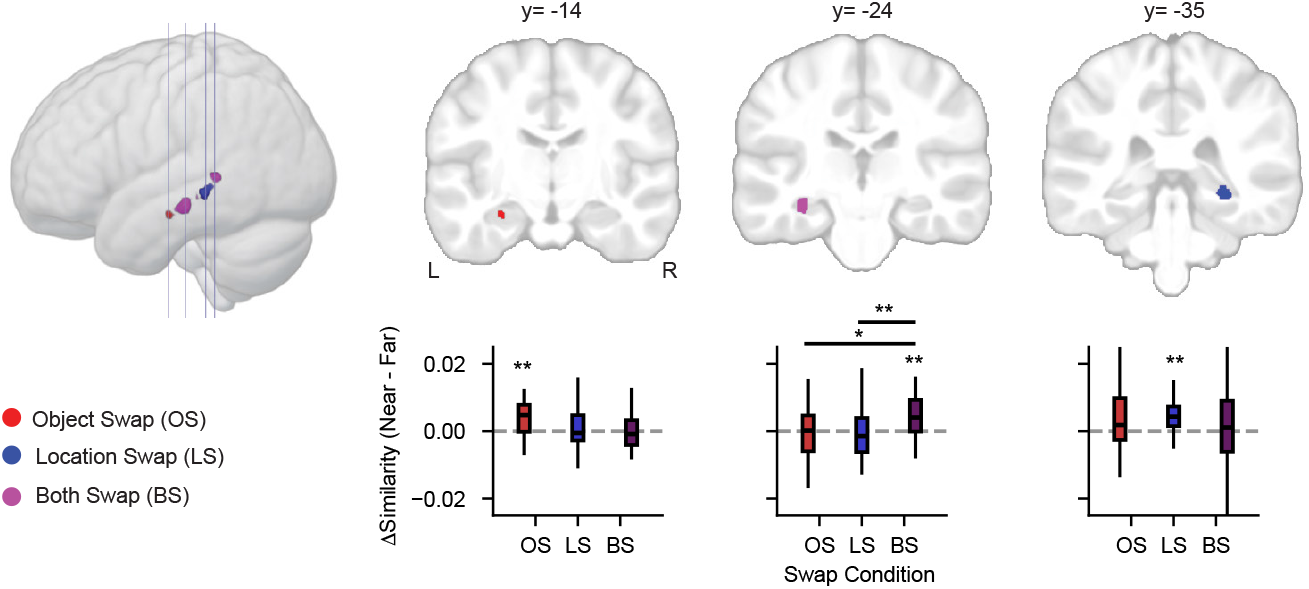
Subregions along the hippocampal long axis show differential sensitivity to semantic and spatial mismatches. Four clusters exhibited significantly greater neural similarity for near vs. far mismatches. Object swaps (OS) showed greater neural similarity for near mismatches in left anterior hippocampus, location swaps (LS) in right posterior hippocampus (pHPC), and both swaps (BS) in left intermediate hippocampus and right pHPC. Significance is indicated as **\*\***, *p <* 0.005; **\***, *p <* 0.05 (one-sample *t*-tests). Box-and-whiskers plots show the distribution of Δ similarity, with the box, line, and whiskers denoting the interquartile range (IQR), the median, and 1.5 times the IQR. See Table S1 for additional details.

### Distinct cortico-hippocampal network dynamics explain functional specialization

Given the presence of subregions with different sensitivity to the spatial or semantic content of sequence mismatches, we next sought to test whether differences in the connectivity of these regions might explain their function. Specifically, we hypothesized that regions sensitive to the semantic content of sequences (aHPC) would be coupled to systems involved in semantic processing such as anterior temporal networks [51] while hippocampal regions involved in detecting visuospatial mismatches (pHPC) would show functional connectivity with early visual and dorsal attention networks [52]. We further reasoned that iHPC may show less specific patterns of connectivity, given its processing of both spatial and semantic attributes.

To test these hypotheses, we completed a seed-based single-trial connectivity analysis [53] looking at changes in the functional connectivity of each mismatch-sensitive subregion (Fig. 5A) during the memory test. As shown in Figure 5B, we observed distinct patterns of connectivity for three mismatch-sensitive subregions. The aHPC subregion connectivity maps revealed regions of the anterior temporal lobe, predominantly in the middle and superior temporal gyrus. Connectivity to the iHPC subregion showed a partially overlapping network of regions, predominantly in the superior temporal gyrus and cingulate cortex. pHPC connectivity maps, on the other hand, showed a markedly different pattern of connectivity with early visual, dorsal parietal, and some areas in the salience and parietal memory networks including the insula and posterior cingulate cortex.

**Figure 5.**
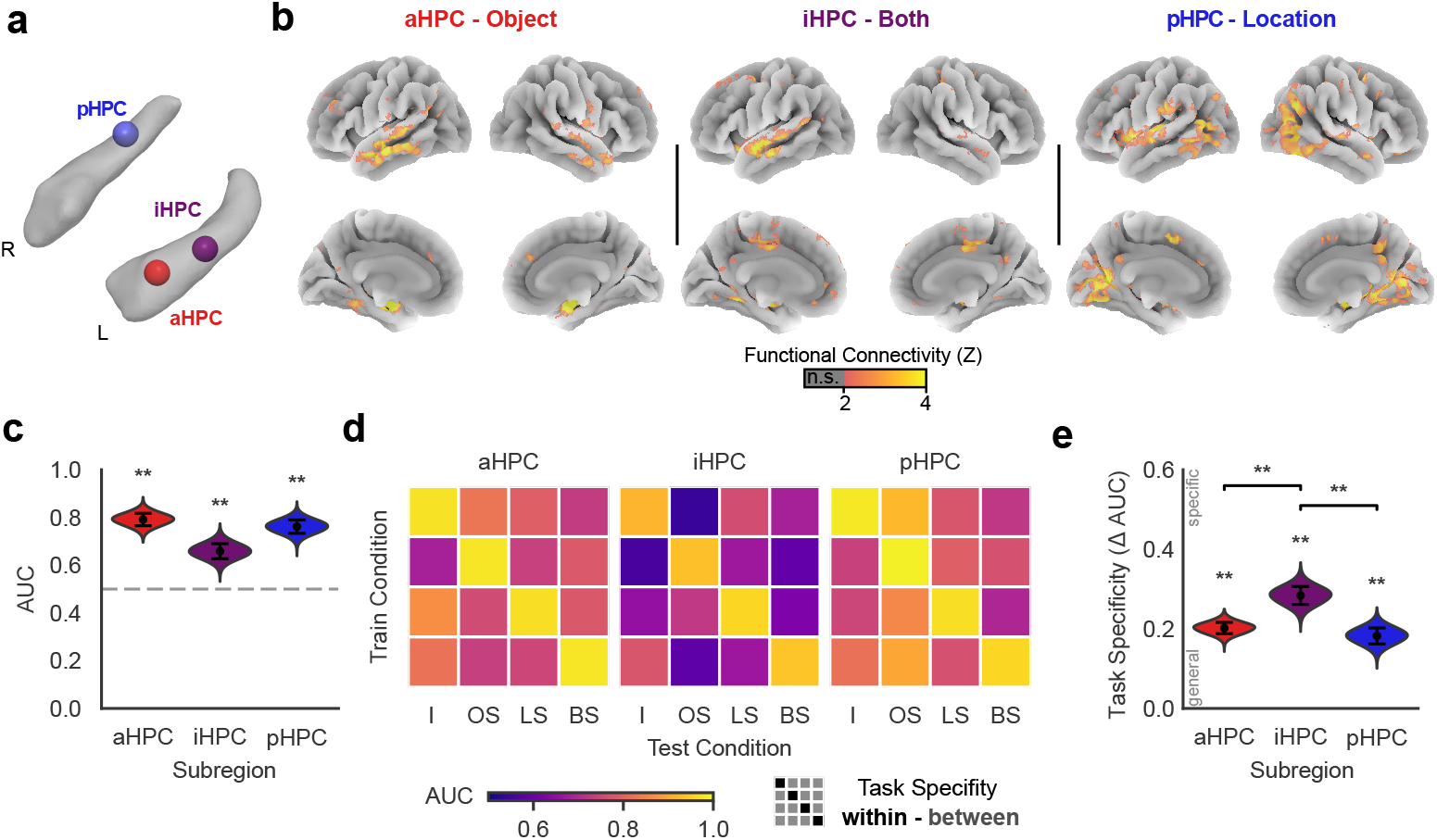
Distinct functional connectivity to the anterior (aHPC), intermediate (iHPC), and posterior (pHPC) hippocampus. (**a**) Locations of seeds within the right (R) and left (L) hippocampus used to generate functional connectivity maps. (**b**) Single-trial connectivity maps reveal distinct functional networks for hippocampal subregions. Maps are thresholded to show regions on the cortical surface with significant functional connectivity with each seed (*p <* 0.05 FDR corrected, permutation test). (**c**) Ridge classifiers successfully discriminate connectivity maps generated from each subregion. Classifier performance is depicted using area under the curve (AUC). **, *p <* 0.005, FDR corrected. Error bars denote bootstrap estimated SEM. (**d**) Generalization testing reveals specificity of hippocampal connectivity. Average AUC across participants (*N* = 28) is depicted for classifiers trained and tested on each combination of intact (I), object swap (OS), location swap (LS), and both swap (BS) trials. (**e**) All functional networks show task specificity, defined as the loss in classifier performance for within vs. between task classification of whole-brain networks (see **d**). Mismatch processing modulates iHPC connectivity to a greater extent than aHPC or pHPC networks. **, *p <* 0.005, FDR corrected, permutation test. See Tables S2-S4 for additional details.

We quantified the distinctiveness of these networks using ridge classification to identify each whole-brain connectivity pattern for a given subregion, excluding all voxels within the hippocampus to prevent local activity from biasing classification performance. As shown in Figure 5C, connectivity patterns were most distinct for aHPC (AUC = 0.79, 95% CI [0.73, 0.84], *p <* 0.001, permutation test), followed by pHPC (AUC = 0.76, 95% CI [0.71, 0.82], *p <* 0.001), with iHPC showing the least discriminability (AUC = 0.66, 95% CI [0.60, 0.72], *p <* 0.001). These results confirm that each mismatch-sensitive subregion is associated with a distinct cortical network.

We next asked whether the connectivity patterns of each hippocampal subregion changed depending on memory demands, or remained stable when different types of information mismatched in memory. To test this, we trained separate classifiers to distinguish connectivity maps for each type of mismatch, and evaluated network specificity by comparing classifier performance when tested within the same versus different task conditions in held-out participants (Fig. 5D).

All three subregions showed significant specificity: aHPC (specificity = 0.20, SEM = 0.01, *p*_FDR_ = 0.001), iHPC (specificity = 0.28, SEM = 0.02, *p*_FDR_ = 0.001), and pHPC (specificity = 0.18, SEM = 0.02, *p*_FDR_ = 0.001). Notably, iHPC showed significantly greater specificity than both aHPC (Δ specificity = 0.08, SEM = 0.03, *p*_FDR_ = 0.003) and pHPC (Δ specificity = 0.10, SEM = 0.03, *p*_FDR_ = 0.003). These findings indicate that connectivity in iHPC changes more readily with memory demands, suggesting greater memory-dependent flexibility compared to aHPC or pHPC.

Together, these results reveal that hippocampal subregions involved in mismatch processing interact with distinct cortical networks that dynamically reconfigure based on the type of representation that mismatches with memory. This functional differentiation supports the idea that long-axis specialization in the hippocampus is shaped by both stable network affiliations and flexible task-dependent interactions.

## Discussion

Using high-resolution fMRI and eye tracking, we found that hippocampal responses during sequence recognition were shaped by the extent to which learned sequences preserved their semantic and spatial structure. When test sequences retained their relational structure, neural similarity varied smoothly along the hippocampal long axis, with higher similarity in anterior and lower similarity in posterior regions. This content-general pattern suggests that gradient-like organization may reflect a general property of hippocampal function when ongoing experience aligns with internal models.

By contrast, when sequence expectations were violated, this long-axis gradient gave way to modular responses localized to discrete hippocampal subregions. The anterior hippocampus was most sensitive to semantic mismatches, the posterior hippocampus to spatial mismatches, and the intermediate hippocampus to combined semantic–spatial violations. These modules exhibited distinct response profiles and cortical coupling patterns, selectively connecting to anterior temporal, visual, and temporo-cingulate networks, respectively. Notably, such reorganization was absent during sequence repetition, suggesting that modular specialization emerges specifically under conditions of prediction error.

These findings reconcile competing models of hippocampal organization by showing that gradient and modular architectures are dynamically recruited depending on memory fidelity and representational demands. Importantly, this framework resonates with evidence from rodent models showing both longitudinal gradients in spatial coding and discrete functional modules aligned with behavioral demands [8, 13]. Yet, the ability to flexibly organize representations by semantic as well as spatial structure may reflect an evolutionarily expanded role for the anterior hippocampus in humans. This anterior expansion co-occurs with connectivity to transmodal cortical areas [23, 24], particularly with networks involved in conceptual abstraction and semantic memory [51, 54]. Notably, comparative work has shown that despite broad conservation of hippocampal structure across primates, the anterior hippocampus in humans exhibits functional reconfiguration aligned with default mode and heteromodal association networks [10]. By leveraging high-resolution fMRI and within-subject functional mapping, our results reveal how this region supports content-specific modules that are anatomically interleaved with, and dynamically recruited in place of, the long-axis gradient under conditions of uncertainty. These findings underscore the importance of studying hippocampal organization in humans, where representational content spans beyond physical space to include abstract, semantic structure.

Our results clarify ongoing debates about hippocampal organization by demonstrating that gradient and modular architectures coexist, but are differentially expressed depending on task demands. Existing models describe a continuous long-axis gradient in precision (i.e., gist to detail), regardless of content type but shaped by neural inputs [8]. Other accounts highlight categorical representational differences across the long axis, such as preferential coding of objects versus scenes in anterior versus posterior regions of the MTL [55–57]. Evidence from rodents also supports modular organization along the long axis, with distinct subregions differing in gene expression [58], lesion effects [59], and intrinsic connectivity [60, 61]. Human studies have tested these frameworks individually [19, 20], but few have examined how both spatial and semantic prediction errors jointly shape hippocampal responses.

By analyzing spatial and semantic mismatches in the same individuals, our findings reveal that gradient-like organization is not a fixed organizational property but a functional mode expressed when input preserves the relational structure of memory. These results challenge models proposing a unitary, continuous gradient and instead suggest that hippocampal coding schemes shift depending on the fidelity of memory-to-input matching, supporting a dual-architecture account in which graded and modular coding flexibly alternate to meet task demands.

Hippocampal activity diverged markedly when spatial and semantic mismatches co-occurred. Although neural similarity followed a smooth anterior–posterior gradient when object identity and location were manipulated independently, this gradient collapsed when both features were altered together. Instead, anterior similarity declined and posterior similarity increased, irrespective of mismatch magnitude. Reduced similarity in aHPC aligns with its proposed role in integrating nonspatial, contextual associations [62], and may reflect disruption of sequence-level representations, given prior evidence linking aHPC to detection of conceptual novelty and temporal structure [33, 63]. This pattern is compatible with a prediction-error mechanism in which hippocampal populations alter their firing patterns to encode violations of expected sequences [64–66]. iHPC and pHPC have been associated with retrieval of temporal context [67], suggesting participants relied more on sequence-level information when both object and location cues alone were unreliable. Increased similarity in pHPC, in turn, may reflect perceptual repetition across study and test phases, consistent with tuning to local visual overlap rather than integrative sequence processing.

The intermediate hippocampus occupies a transitional zone along the long axis, serving as a critical test as to whether hippocampal organization reflects a smooth gradient or discretely specialized subregions. In our data, iHPC did not exhibit a monotonic shift between aHPC and pHPC functions. Instead, its pattern resembled a convergence of anterior and posterior responses, supporting integrative representations. This aligns with rodent evidence that iHPC encodes reward location and integrates broad motivational context with spatial detail [68, 69]. Although the anterior hippocampus is often linked to affective and conceptual processing [21, 70], and posterior regions to detailed visuospatial encoding [12], the iHPC appears uniquely positioned to process co-occurring semantic and spatial features. Our findings support the view that intermediate regions within the human hippocampal body play a distinct computational role in combining representational streams.

As predicted, we found that aHPC and pHPC are functionally embedded in distinct large-scale cortical systems during semantic and spatial memory, respectively. The aHPC network primarily included anterior and ventral temporal regions, consistent with proposals wherein anterior temporal systems support semantic and item-based processing [51, 71, 72]. Connectivity with these regions reflects the anterior hippocampus’s strong coupling with heteromodal association areas, in line with prior work linking aHPC to semantic abstraction and high-level cognition [73, 74]. In contrast, pHPC was more strongly connected to early visual cortex, dorsal parietal areas, and posterior cingulate which are associated with perceptual detail, spatial attention, and memory-guided orienting [22, 23, 28]. These findings extend prior resting-state studies [28, 29], revealing that long-axis differences in cortical coupling are also engaged during active, content-specific memory processing. They further raise the question as to how iHPC functionally connects to these systems, potentially with overlapping connectivity with either pole.

The intermediate hippocampus exhibited a distinct connectivity profile involving lateral temporal cortex regions that were posterior to those connected with aHPC. This topographic shift mirrors the pattern of interdigitated cortical systems along the anterior-posterior axis [75], and is consistent with the idea that iHPC supports integration of semantic and spatial information across cognitive states. These distinctions also parallel dorsal–ventral organizational schemes in rodents, where dorsal hippocampus supports fine-grained spatial coding and ventral regions contribute to integrative and contextual representations [21]. Rodent studies further suggest that iHPC flexibly combines signals from both poles to guide behavior under uncertainty [69]. Together, these findings suggest that anterior, intermediate, and posterior hippocampal subregions are embedded in dissociable large-scale systems, with functional connectivity patterns shaped by both representational content and task demands.

Our study is not without potential limitations. For example, we restricted our sampling of semantic and spatial information. Our stimulus set captures only a fraction of the semantic richness found in everyday experience, and the spatial manipulations offer a simplified model of real-world spatial variation. However, the stimuli were specifically designed to provide a broad and diverse sampling of object concepts grounded in human behavioral norms and visual representations [38, 76, 77]. Prior studies have shown that even simplified or highly controlled stimulus sets can robustly engage semantic memory networks [78, 79] and reveal stable representational patterns in the hippocampus and cortex [47, 80]. Likewise, controlled spatial paradigms including virtual environments and simplified visuospatial arrays have reliably elicited hippocampal activity related to spatial navigation [5, 81] and visuospatial organization [18, 82, 83]. Such findings combined with the strong behavioral effects observed in our task validate our paradigm.

One key limitation of the current study is the inability to fully capture the timing and causal mechanisms of hippocampal memory processing. Although fMRI offers valuable spatial precision and whole-brain coverage, it lacks the temporal resolution to detect rapid neural computations that occur within hundreds of milliseconds. Intracranial recordings have shown rapid mismatch detection is coordinated by hippocampal theta oscillations [37, 84], revealing temporal dynamics inaccessible to fMRI. Additionally, most human evidence remains correlational. While lesion studies demonstrate the hippocampus’s critical role in memory [85, 86], selective damage to anterior, posterior, or intermediate subregions is rare, limiting causal inference about regional specialization. Moving forward, combining stimulation-based methods [87] with high-temporal-resolution recordings will be essential to precisely delineate the fast, region-specific mechanisms that support hippocampal memory processing and to extend the insights gained from this work.

## Conclusion

By jointly manipulating spatial and semantic memory, we reveal how hippocampal organization flexibly adapts to cognitive demands. When input aligns with learned sequences, neural similarity follows a smooth gradient along the long axis. But when memory is violated, modular subregions emerge with content-specific tuning and distinct connectivity profiles. These findings suggest that hippocampal coding schemes are not fixed but reflect adaptive computations that balance prediction, error detection, and representational precision depending on the content and strength of memory.

## Materials and Methods

### Participants

Thirty-four human participants (18 female; on average 26, range 18–49 years of age) were recruited for the study. Participants were compensated $30 per hour for their time upon study completion. All participants had normal or corrected-to-normal vision, no neuropsychiatric history, no psychotropic drug use, and no MRI contraindications. Data from 6 participants were excluded from analysis due to task performance at or below chance for any condition. The research protocol was approved by the University of Chicago Biological Sciences Division Institutional Review Board, and all participants provided informed written consent.

### Experimental procedure

The study began with a short behavioral training session, during which participants were instructed on the task, scanning protocol, and eye-tracking procedures. Participants performed a practice version of the task with four example trials and were provided feedback on their performance. Two fMRI sessions followed this training session, one immediately after and one typically 2 (median; range 1–38) days following the first session. Repeated imaging sessions provided additional data for precision mapping of hippocampal gradients.

During each fMRI session, participants underwent eight functional runs while completing a sequence mismatch detection task. The stimuli were presented using a Psychology Software Tools, Inc. Hyperion projector on a 60.0 cm (width) x 45.0 cm (height) screen with a resolution of 1024 x 768 pixels at 60 Hz. Participants viewed the display through a mirror mounted on the RF coil 12 cm from the eye, with an additional 127 cm from the mirror to screen. Stimuli were presented on a circular grid placed in the center of the display with a diameter of 18.39^*°*^ (768 pixels). The ability of participants to see the entire display area was confirmed prior to each session. Psychophysics Toolbox (version 3.0.18; 88, 89) was run from MATLAB (Natick, MA) on a Windows 10 control PC to display the task within the scanner bore.

Throughout the study, participants learned 256 total object sequences divided into 16 blocks of 16 trials each. During study, each object in the sequence was presented on screen for 750 ms with an inter-stimulus interval of 250 ms, for a total of 5 seconds. Individual objects spanned 3.54^*°*^ (150 pixels) and were displayed at one of eight locations at a distance of 6.97^*°*^ (290 pixels) from the center of a fixation grid that persisted for the duration of the trial. The grid was included to facilitate fixation when no objects were on the screen [90, 91], and to provide a spatial reference for the location of each object. To alert participants to an upcoming sequence, a red dot was presented at the center of the screen up to 1 s prior to object presentation. Following a 14-second delay, a test sequence with the same timing as the original study sequence was presented. Following the test sequence, there was a 3–5 second delay prior to a response period, jittered to allow separation of response-related activity. The inter-trial interval was also jittered between 8–10 seconds.

Objects in each sequence were drawn from a subset of unique, real-life manmade and natural objects from the THINGS dataset [38]. Exclusions were made based on object memorability (corrected recognition below 65%, 92), emotional arousal (the top and bottom 5 percent of objects ranked on average arousal ratings, 92), and pleasantness (negative ratings below 4 on a Likert scale of 1–7, 92). Additionally, objects highly associated with sexual desire were excluded (the top 5 percent based on a neural network model of emotion, 93) in addition to eight additional objects based on subjective judgment of the authors. These exclusion criteria were used to ensure objects were memorable but relatively neutral in content and arousal.

The spatial and semantic content of objects within each sequence was carefully controlled. At study, temporally adjacent objects in each sequence were either near or far based on their similarity in spatial and semantic dimensions.

Semantic similarity was derived from a semantic embedding of objects in a neural network model of human similarity judgments on a triplet odd-one-out task [38]. The similarity between two objects was computed from their 66-dimensional vector representations:

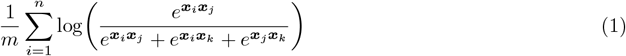

where ***x*** corresponds to the vector representation of each of *m* objects, based on each triplet with items *i, j*, and *k* from the full set of *n* triplets. Two objects were considered near one another if their semantic similarity was greater than 0.6 and far from one another if their semantic similarity was less than 0.35. For each sequence, we also made sure that each object was distant from all other non-adjacent objects in the sequence (semantic similarity less than 0.35).

Spatial similarity was determined based on the distance between consecutively presented objects. From the eight possible locations distributed uniformly around the background fixation grid (45^*°*^ spacing, 15^*°*^ offset from the axis of the screen, see Fig. 1), near locations were in adjacent positions (*±* 45^*°*^) and far locations were similarly spaced across the circle (135^*°*^ – 225^*°*^). Spatial locations were never repeated within a sequence, and all locations were equally viewed across each functional run and session.

We used the controlled structure of each sequence to test memory for semantic and spatial content at different scales. In a 2 *×* 2 design, we swapped either the object or location of the final two stimuli in the sequence. Thus, a given test sequence was either an intact, object swap, location swap, or both swap trial. Half the trials started with near and half with far transitions, allowing us to examine the effects of distance on memory and hippocampal activity. During test, participants were tasked with detecting sequence mismatches. During the response period, participants judged whether the probe sequence differed from the study sequence in any way (judging the sequence as old or new). Their responses were recorded using a 5-button response box affixed to the participant’s right hand.

### Eye tracking

Eye movements were recorded at 500 Hz using an EyeLink 1000 Plus remote tracking system (software version 5.16, SR Research Ltd., Ontario, Canada). The eye-tracking camera apparatus was placed roughly 91 cm away from the participant’s eyes at the head of the scanner bore. Each participant confirmed that they had an unobstructed view of the entire screen before beginning the experiment. A five-point gaze calibration and validation were performed at the beginning of each task block. The average validation error across all participants was 0.44 degrees of visual angle (29 participants, *SD* = 0.18). Continuous eye-tracking data were parsed into fixation, saccade, and blink events. Motion (0.15°), velocity (30^*°*^*/s*), and acceleration (8000^*°*^*/s*^2^) thresholds determined saccade events. Blinks were determined by pupil size, and remaining epochs below detection thresholds were classified as fixations. The location of each fixation was computed as the average gaze position throughout the period of fixation.

We focused on two eye-tracking measures to examine spatial and object memory. To assess spatial memory, we measured the total number of fixations made just prior to the critical fourth stimulus at test. We reasoned that fixations to the original object location (from study) reflected spatial memory, which we refer to as predictive fixations. We labeled fixations as predictive if they occurred between 500 and 100 ms before the fourth object at test within the actual boundaries of the object location. To assess whether these fixations reflected spatial memory, we compared the number of predictive fixations to fixations to all other possible object locations. We additionally assessed spatial mismatches by comparing the number of predictive fixations to fixations to the mismatched location on LS and BS trials. To assess semantic memory for objects in the sequence, we measured the duration of the first fixation made to objects in a given sequence. First fixation durations were quantified by measuring the length of time a participant first spent fixating on the critical fourth image in the sequence at test.

### MRI acquisition

All fMRI data were collected at the University of Chicago MRI Research Center on a Philips Ingenia 3 Tesla scanner. Structural images were acquired using a T1-weighted 3D magnetization-prepared rapid gradient-echo sequence (field of view = 256 mm, echo time (TE) = 3 ms, repetition time (TR) = 7 ms, flip angle = 8°, voxel resolution: 1.0 *×* 1.0 *×* 1.0 mm). Functional images were acquired with a MultiBand 3 echo planar imaging sequence (TE = 20 ms, TR = 2 s, flip angle = 80°, matrix size = 128 *×* 128, slice thickness = 1.8 mm, field of view = 220 mm, 81 slices, voxel resolution = 1.625 *×* 1.625*×* 1.8 mm).

### fMRI preprocessing

Results included in this manuscript come from preprocessing performed using *fMRIPrep* 23.1.4 [94]. See Supplementary Methods for details.

### Hippocampal unfolding and gradient estimation

T1-weighted structural images were additionally processed with HippUnfold [49, 50] which registers hippocampal volumes to an unfolded surface space and performs subfield segmentation. A Laplace gradient is calculated between the anterior and posterior ends of the hippocampus (for details, see 48). This gradient falls within hippocampal grey matter and provides native space Laplace coordinates that account for variability in hippocampal curvature. Coordinates in the AP direction were used in all regression analyses looking at representational change along the long axis.

### General linear modeling

For each functional run, single-trial estimates of neural responses were generated using a general linear model (GLM) in *Nipype*. The study and test periods of each trial were modeled by convolving the canonical hemodynamic response function in SPM (version 12) with a 5s block. The response period was modeled in the same fashion but with a 3s duration. A set of nuisance regressors was included in the model to account for head motion and physiological noise. These included framewise displacement, 6 motion parameters (x, y, z, roll, pitch, and yaw), and the top 6 anatomical component-based correction parameters (aCompCor from *fMRIPrep*). We fitted a least-squares-all model to estimate beta coefficients for subsequent analyses [95]. Modeling each trial separately allowed for the precise examination of trial-level neural signals while accounting for confounds related to head motion and physiological noise.

### Representational similarity analysis

We tested for the presence of a functional gradient along the hippocampal long axis using representational similarity analysis (RSA) on neural activity patterns derived from the general linear model described above. RSAs were computed separately for each condition and swap distance. We implemented a search-light analysis where spheres with a 4 mm radius were centered on each hippocampal voxel. Spheres near the boundaries of the hippocampus were masked to include only those voxels within the hippocampus. For each sphere, we extracted multi-voxel activity patterns from unsmoothed beta images for each trial, resulting in a trial-by-voxel activation matrix. We computed the average pairwise Pearson correlation between trials within each condition, Fisher Z-transforming the values prior to averaging. To assess the effects of mismatch strength, we computed neural similarity differences between near and far trials separately for each swap type.

To examine changes in neural similarity along the hippocampal long axis, we estimated repeated-measures correlations [96] between the anterior–posterior (AP) coordinate and neural similarity for each condition. These correlations were computed across searchlight centers for each subject and condition. We then compared the resulting AP-similarity correlations between repeat and swap trials, and between near and far swaps, to assess how gradient strength was modulated by mismatch strength and content.

### Functional connectivity and classification of hippocampal networks

We assessed whether anterior, intermediate, and posterior hippocampal subregions were embedded in functionally distinct networks using condition-level multivariate classification of connectivity maps. Seed regions were defined from mismatch-sensitive clusters along the hippocampal long axis (see Fig. 5b and Table S1). For each subject, a whole-brain functional connectivity map was computed for each seed and condition, resulting in one map per seed–condition pair.

Classification was performed using ridge regression classifiers. Before model training, input features were standardized to zero mean and unit variance with principal component analysis applied for feature reduction, retaining 100 components. This standardization and dimension reduction were performed using only the training data in each fold of a leave-one-subject-out cross-validation procedure. The same parameters were applied to the held-out test data.

One-vs-all classification performance was quantified using area under the ROC curve, computed from decision function scores. AUC scores were pooled across folds and computed separately for each connectivity profile. To assess task-specific network reconfiguration, we performed cross-condition generalization tests. For each condition, a classifier was trained to predict network identity using functional connectivity maps from that condition and tested on maps from all possible conditions. Network specificity was defined as the difference in AUC between within-condition and between-condition performance.

### Statistical procedures

Univariate comparisons of behavioral accuracy and eye movement measures were tested using one-sample or paired-sample t-tests. Effect sizes were quantified using Hedges’ g [97].

We used permutation procedures to assess the significance of long-axis gradients in neural similarity. For each condition, we evaluated the significance of repeated-measures correlation coefficients using a sign-flipping procedure across subjects to generate a null distribution (n = 5000). To test differences in gradient strength between near and far trials, we permuted the condition labels within subjects and computed a surrogate distribution of difference scores. Similar permutation methods were used to test for differences in classifier performance and network specificity, randomly permuting the network labels within subjects. For these tests, FDR-correction for multiple comparisons was applied.

## Acknowledgments

We would like to thank the University of Chicago’s MRI Research Core and Research Computing Center for their support in the acquisition and analysis of imaging data. This research was supported by the National Institutes of Health grant R01MH128552.

## Supplementary Information

### Supplementary Methods

#### Anatomical data preprocessing

Anatomical data were preprocessed using *fMRIPrep* 23.1.4 [98]. A total of 2 T1-weighted (T1w) images were found per subject within the input BIDS dataset. All of them were corrected for intensity non-uniformity (INU) with `N4BiasFieldCorrection` [99], distributed with ANTs [100]. The T1w-reference was then skull-stripped with a *Nipype* implementation of the `antsBrainExtraction.sh` workflow (from ANTs), using OASIS30ANTs as target template. Brain tissue segmentation of cerebrospinal fluid (CSF), white-matter (WM) and gray-matter (GM) was performed on the brain-extracted T1w using fast (FSL version 6.0.7.7, [101]). An anatomical T1w-reference map was computed after registration of 2 T1w images (after INU-correction) using `mri_robust_template` (FreeSurfer version 7.3.2, [102]). Brain surfaces were reconstructed using `recon-all` (FreeSurfer version 7.3.2, [103]), and the brain mask estimated previously was refined with a custom variation of the method to reconcile ANTs-derived and FreeSurfer-derived segmentations of the cortical gray-matter of Mindboggle [104]. Volume-based spatial normalization to one standard space (MNI152NLin2009cAsym) was performed through nonlinear registration with antsRegistration (ANTs, version 2.5.1), using brain-extracted versions of both T1w reference and the T1w template. The following template was selected for spatial normalization and accessed with *Template-Flow* (version 23.0.0, [105]): *ICBM 152 Nonlinear Asymmetrical template version 2009c* [106].

#### Functional data preprocessing

For each of the BOLD runs found per subject (up to 16 across sessions), the following preprocessing was performed. First, a reference volume and its skull-stripped version were generated using a custom methodology of *fMRIPrep*. Head-motion parameters with respect to the BOLD reference (transformation matrices, and six corresponding rotation and translation parameters) are estimated before any spatiotemporal filtering using `mcflirt` (FSL, [107]). BOLD runs were slice-time corrected to 1.0s (0.5 of slice acquisition range 0s-2.0s) using `3dTshift` from AFNI [108]. The BOLD time-series (including slice-timing correction when applied) were resampled onto their original, native space by applying the transforms to correct for head motion. These resampled BOLD time-series will be referred to as preprocessed BOLD in original space, or just preprocessed BOLD. The BOLD reference was then co-registered to the T1w reference using `bbregister` (FreeSurfer) which implements boundary-based registration [109]. Co-registration was configured with six degrees of freedom. Several confounding time-series were calculated based on the pre-processed BOLD: framewise displacement (FD), DVARS and three region-wise global signals. FD was computed using two formulations following Power (absolute sum of relative motions, [110]) and Jenkinson (relative root mean square displacement between affines, [107]). FD and DVARS are calculated for each functional run, both using their implementations in *Nipype* following the definitions Power et al. [110]. The three global signals are extracted within the CSF, the WM, and the whole-brain masks. Additionally, a set of physiological regressors were extracted to allow for component-based noise correction (*CompCor*, [111]). Principal components are estimated after high-pass filtering the preprocessed BOLD time-series (using a discrete cosine filter with 128s cut-off) for the two *CompCor* variants: temporal (tCompCor) and anatomical (aCompCor). tCompCor components are then calculated from the top 2% variable voxels within the brain mask. For aCompCor, three probabilistic masks (CSF, WM and combined CSF+WM) are generated in anatomical space. The implementation differs from that of Behzadi et al. in that instead of eroding the masks by 2 pixels on BOLD space, a mask of pixels that likely contain a volume fraction of GM is subtracted from the aCompCor masks. This mask is obtained by dilating a GM mask extracted from the FreeSurfer’s *aseg* segmentation, and it ensures components are not extracted from voxels containing a minimal fraction of GM. Finally, these masks are resampled into BOLD space and binarized by thresholding at 0.99 (as in the original implementation). Components are also calculated separately within the WM and CSF masks. For each CompCor decomposition, the *k* components with the largest singular values are retained, such that the retained components’ time series are sufficient to explain 50 percent of variance across the nuisance mask (CSF, WM, combined, or temporal). The remaining components are dropped from consideration. The head-motion estimates calculated in the correction step were also placed within the corresponding confounds file. The confound time series derived from head motion estimates and global signals were expanded with the inclusion of temporal derivatives and quadratic terms for each [112]. Frames that exceeded a threshold of 0.5 mm FD or 1.5 standardized DVARS were annotated as motion outliers. Additional nuisance timeseries are calculated by means of principal components analysis of the signal found within a thin band (*crown*) of voxels around the edge of the brain, as proposed by Patriat et al. [113]. The BOLD time-series were resampled into standard space, generating a preprocessed BOLD run in MNI152NLin2009cAsym space. First, a reference volume and its skull-stripped version were generated using a custom methodology of *fMRIPrep*. All resamplings can be performed with a single interpolation step by composing all the pertinent transformations (i.e. head-motion transform matrices, susceptibility distortion correction when available, and co-registrations to anatomical and output spaces). Gridded (volumetric) resamplings were performed using `antsApplyTransforms` (ANTs), configured with Lanczos interpolation to minimize the smoothing effects of other kernels [114]. Non-gridded (surface) resamplings were performed using `mri_vol2surf` (FreeSurfer).

**Table S1.**
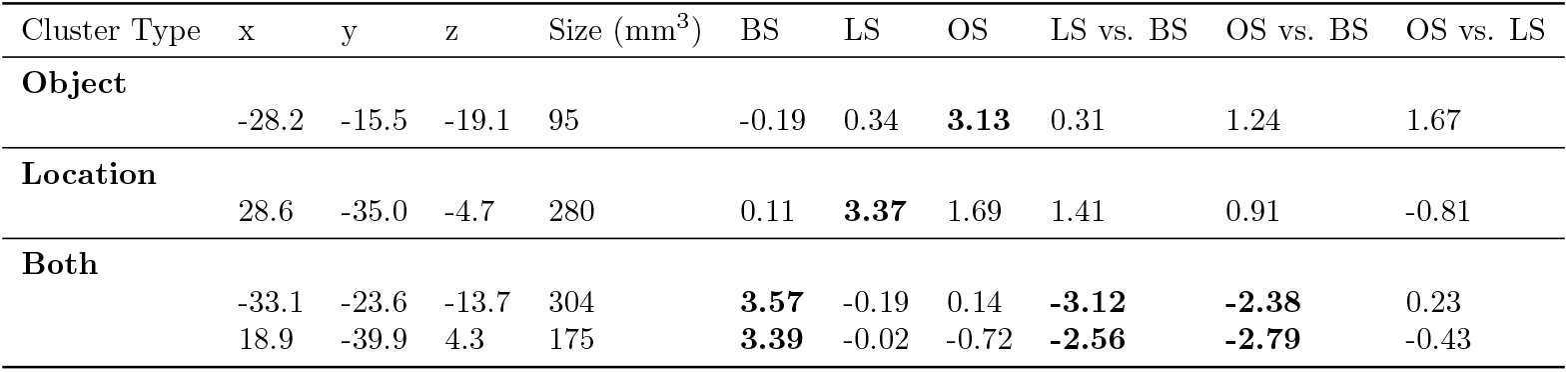
Searchlight clusters showing mismatch sensitivity for different memory content. Coordinates (x, y, z) correspond to the voxel with the maximal *t*-value within each cluster extent. Statistics reflect one-sample *t*-tests of mismatch sensitivity (ΔRSA, near vs. far) against zero for each condition: both swap (BS), location swap (LS), and object swap (OS). Paired *t*-tests indicate direct comparisons between mismatch types. Post-hoc significance is indicated in **boldface** (*p <* .05, FDR-corrected across all post-hoc tests).

**Table S2.** Clusters of significant functional connectivity were identified for the anterior hippocampus (aHPC), with an FDR-corrected threshold (*p <* 0.05, *t >* 2.93), a minimum cluster size of 100 voxels, and a minimum peak-to-peak distance of 20 mm. For each cluster, the peak voxel (highest *t*-statistic) was extracted and labeled using the Harvard-Oxford subcortical and cortical structural atlases [115–119]. MTG, middle temporal gyrus. STG, superior temporal gyrus. SMG, supramarginal gyrus.

| Seed | Cluster | x | y | z | Hemisphere | *t* | Size (mm<sup>3</sup>) | Region |
| --- | --- | --- | --- | --- | --- | --- | --- | --- |
| aHPC | 1 | -26.6 | -13.9 | -19.1 | L | 9.0 | 31536 | Hippocampus |
|  | 1a | -65.6 | -44.8 | 6.1 | L | 4.2 |  | MTG, temporooccipital part |
|  | 1b | -46.1 | 15.4 | -19.1 | L | 4.1 |  | Temporal Pole |
|  | 1c | -67.2 | -33.4 | 4.3 | L | 4.1 |  | STG, posterior division |
|  | 2 | 28.6 | -12.2 | -19.1 | R | 5.6 | 8588 | Hippocampus |
|  | 3 | 56.2 | 12.1 | -19.1 | R | 5.0 | 5827 | Temporal Pole |
|  | 3a | 70.9 | -35.0 | 7.9 | R | 4.2 |  | MTG, posterior division |
|  | 4 | 61.1 | -5.8 | -26.3 | R | 4.9 | 1487 | MTG, anterior division |
|  | 5 | -55.9 | -52.9 | 20.5 | L | 4.8 | 2932 | Angular Gyrus |
|  | 6 | 61.1 | -5.8 | 11.5 | R | 4.4 | 1259 | Central Opercular Cortex |
|  | 7 | -3.9 | 34.9 | 24.1 | L | 4.3 | 484 | Paracingulate Gyrus |
|  | 8 | 9.1 | -69.1 | -15.5 | R | 4.2 | 1839 | Lingual Gyrus |
|  | 9 | 62.8 | -4.1 | 24.1 | R | 4.1 | 1026 | Precentral Gyrus |
|  | 10 | -41.2 | 36.5 | 34.9 | L | 3.9 | 1273 | Middle Frontal Gyrus |
|  | 11 | 36.8 | -0.9 | 13.3 | R | 3.8 | 551 | Insular Cortex |
|  | 12 | 9.1 | 65.8 | 25.9 | R | 3.8 | 608 | Frontal Pole |
|  | 13 | -44.5 | -41.5 | 42.1 | L | 3.7 | 789 | SMG, posterior division |
|  | 14 | 15.6 | 46.2 | 38.5 | R | 3.7 | 632 | Frontal Pole |
|  | 15 | -62.4 | -7.4 | 40.3 | L | 3.7 | 903 | Postcentral Gyrus |
|  | 16 | 2.6 | 39.8 | 27.7 | R | 3.6 | 527 | Paracingulate Gyrus |
|  | 17 | 10.8 | -69.1 | 22.3 | R | 3.6 | 480 | Cuneal Cortex |
|  | 18 | -26.6 | 20.2 | -22.7 | L | 3.5 | 708 | Frontal Orbital Cortex |
|  | 19 | -10.4 | 62.5 | 24.1 | L | 3.4 | 494 | Frontal Pole |

**Table S3.** Clusters of significant functional connectivity were identified for the intermediate hippocampus (iHPC), with an FDR-corrected threshold (*p <* 0.05, *t >* 2.86), a minimum cluster size of 100 voxels, and a minimum peak-to-peak distance of 20 mm. For each cluster, the peak voxel (highest *t*-statistic) was extracted and labeled using the Harvard-Oxford subcortical and cortical structural atlases [115–119]. MTG, middle temporal gyrus. STG, superior temporal gyrus. SMG, supramarginal gyrus.

| Seed | Cluster | x | y | z | Hemisphere | *t* | Size (mm<sup>3</sup>) | Region |
| --- | --- | --- | --- | --- | --- | --- | --- | --- |
| iHPC | 1 | -33.1 | -23.6 | -13.7 | L | 8.6 | 11540 | Hippocampus |
|  | 1a | 7.5 | -56.1 | -11.9 | R | 3.6 |  | Cerebellum |
|  | 2 | -55.9 | -4.1 | -13.7 | L | 5.8 | 15680 | MTG, anterior division |
|  | 2a | -59.1 | -48.0 | 16.9 | L | 4.1 |  | SMG, posterior division |
|  | 2b | -49.4 | 10.5 | -8.3 | L | 4.0 |  | Temporal Pole |
|  | 2c | -57.5 | -43.1 | 20.5 | L | 3.5 |  | SMG, posterior division |
|  | 3 | 20.5 | -7.4 | -10.1 | R | 5.5 | 3555 | Amygdala |
|  | 4 | -28.2 | -41.5 | 56.5 | L | 4.9 | 14420 | Superior Parietal Lobule |
|  | 4a | -3.9 | 39.8 | 49.3 | L | 3.9 |  | Superior Frontal Gyrus |
|  | 4b | -52.6 | -23.6 | 52.9 | L | 3.9 |  | Postcentral Gyrus |
|  | 4c | -3.9 | 49.5 | 47.5 | L | 3.9 |  | Frontal Pole |
|  | 5 | 10.8 | -25.2 | 47.5 | R | 4.7 | 12153 | Precentral Gyrus |
|  | 6 | -3.9 | 56.0 | -6.5 | L | 4.6 | 560 | Frontal Pole |
|  | 7 | 56.2 | -43.1 | 4.3 | R | 4.6 | 983 | MTG, temporooccipital part |
|  | 8 | 22.1 | -46.4 | 67.3 | R | 4.5 | 13579 | Superior Parietal Lobule |
|  | 8a | 59.5 | -17.1 | 49.3 | R | 3.8 |  | Postcentral Gyrus |
|  | 9 | -18.5 | -25.2 | 76.3 | L | 4.0 | 860 | Precentral Gyrus |
|  | 10 | 5.9 | 52.8 | 45.7 | R | 4.0 | 674 | Frontal Pole |
|  | 11 | -51.0 | -41.5 | 56.5 | L | 3.7 | 727 | SMG, posterior division |
|  | 12 | 15.6 | -31.8 | 61.9 | R | 3.7 | 537 | Precentral Gyrus |
|  | 13 | 67.6 | -17.1 | 0.7 | R | 3.5 | 817 | STG, posterior division |

**Table S4.** Clusters of significant functional connectivity were identified for the posterior hippocampus (pHPC), with an FDR-corrected threshold (*p <* 0.05, *t >* 2.39), a minimum cluster size of 100 voxels, and a minimum peak-to-peak distance of 20 mm. For each cluster, the peak voxel (highest *t*-statistic) was extracted and labeled using the Harvard-Oxford subcortical and cortical structural atlases [115–119]. SMA, supplemental motor area. TFC, temporo-fusiform cortex.

| Seed | Cluster | x | y | z | Hemisphere | *t* | Size (mm<sup>3</sup>) | Region |
| --- | --- | --- | --- | --- | --- | --- | --- | --- |
| pHPC | 1 | 30.2 | -35.0 | -4.7 | R | 8.9 | 285667 | Hippocampus |
|  | 1a | -13.6 | -64.2 | 4.3 | L | 5.6 |  | Intracalcarine Cortex |
|  | 1b | 15.6 | -2.5 | -11.9 | R | 5.6 |  | Cerebellum |
|  | 1c | -10.4 | -64.2 | 0.7 | L | 5.5 |  | Lingual Gyrus |
|  | 2 | 54.6 | 8.9 | -17.3 | R | 4.6 | 6416 | Temporal Pole |
|  | 3 | -5.5 | 5.6 | 49.3 | L | 4.5 | 2281 | SMA |
|  | 4 | 27.0 | 41.4 | -17.3 | R | 4.5 | 3327 | Frontal Pole |
|  | 5 | 9.1 | 4.0 | 52.9 | R | 4.4 | 4681 | SMA |
|  | 5a | 2.6 | 54.4 | 42.1 | R | 4.0 |  | Superior Frontal Gyrus |
|  | 6 | -12.0 | 44.6 | 43.9 | L | 4.3 | 1216 | Frontal Pole |
|  | 7 | 43.2 | -51.2 | -51.5 | R | 4.1 | 9558 | Cerebellum |
|  | 8 | -33.1 | -44.8 | 56.5 | L | 4.1 | 2813 | Superior Parietal Lobule |
|  | 9 | 23.8 | 10.5 | -40.7 | R | 3.9 | 1459 | Temporal Pole |
|  | 10 | 54.6 | 39.8 | -1.1 | R | 3.8 | 1278 | Frontal Pole |
|  | 11 | -41.2 | 43.0 | -1.1 | L | 3.7 | 2542 | Frontal Pole |
|  | 12 | -7.1 | 36.5 | 22.3 | L | 3.5 | 1197 | Cingulate Gyrus, anterior division |
|  | 13 | -3.9 | 10.5 | 27.7 | L | 3.5 | 608 | Cerebral White Matter |
|  | 14 | 53.0 | -23.6 | 9.7 | R | 3.5 | 836 | Planum Temporale |
|  | 15 | -26.6 | 0.8 | 52.9 | L | 3.5 | 2752 | Middle Frontal Gyrus |
|  | 16 | -26.6 | -52.9 | 69.1 | L | 3.4 | 541 | Superior Parietal Lobule |
|  | 17 | -8.8 | -49.6 | 60.1 | L | 3.3 | 798 | Precuneous Cortex |
|  | 18 | -42.9 | 25.1 | 45.7 | L | 3.3 | 708 | Middle Frontal Gyrus |
|  | 19 | 33.5 | -46.4 | -35.3 | R | 3.2 | 584 | Cerebellum |
|  | 20 | -38.0 | -7.4 | -44.3 | L | 2.9 | 1040 | TFC, anterior division |

## Supplementary Movies

All movies show example trials from a single participant with simultaneously recorded eye movements. The current gaze position is indicated in blue.

**Movie S1**. Example trial with an intact sequence.

**Movie S2**. Example trial with an object swap.

**Movie S3**. Example trial with a location swap.

**Movie S4**. Example trial with both an object and location swap.

## Notes

### Competing Interest Statement

The authors have declared no competing interest.

